# Integrated Genomics Approaches Identify Transcriptional Mediators and Epigenetic Responses to Afghan Desert Particulate Matter in Small Airway Epithelial Cells

**DOI:** 10.1101/2022.04.27.489761

**Authors:** Arnav Gupta, Sarah K. Sasse, Reena Berman, Margaret A. Gruca, Robin D. Dowell, Hong Wei Chu, Gregory P. Downey, Anthony N. Gerber

## Abstract

Military Deployment to Southwest Asia and Afghanistan and exposure to toxic airborne particulates has been associated with an increased risk of developing respiratory disease, collectively termed deployment-related respiratory diseases (DRRD). Our knowledge about how particulates mediate respiratory disease is limited, precluding the appropriate recognition or management. Central to this limitation is the lack of understanding of how exposures translate into dysregulated cell identity with dysregulated transcriptional programs. The small airway epithelium is involved in both the pathobiology of DRRD and fine particulate matter deposition. To characterize small airway epithelial cell epigenetic and transcriptional responses to Afghan desert particulate matter (APM) and investigate the functional interactions of transcription factors that mediate these responses, we applied two genomics assays, the assay for transposase accessible chromatin with sequencing (ATAC-seq) and Precision Run-on sequencing (PRO-seq). We identified activity changes in a series of transcriptional pathways as candidate regulators of susceptibility to subsequent insults, including signal-dependent pathways, such as loss of cytochrome P450 or P53/P63, and lineage-determining transcription factors, such as GRHL2 loss or TEAD3 activation. We further demonstrated that TEAD3 activation was unique to APM exposure despite similar inflammatory responses when compared with wood smoke particle exposure and that P53/P63 program loss was uniquely positioned at the intersection of signal-dependent and lineage-determining transcriptional programs. Our results establish the utility of an integrated genomics approach in characterizing responses to exposures and identify genomic targets for the advanced investigation of the pathogenesis of DRRD.

## Introduction

The development of respiratory disease following post-9/11 deployment to Southwest Asia (SW Asia) and Afghanistan and exposure to associated airborne hazards, collectively termed deployment-related respiratory diseases (DRRD), is an important clinical problem that has had a major adverse impact on the long-term health of Veterans. DRRD encompasses several respiratory conditions, such as asthma, constrictive bronchiolitis and chronic rhinosinusitis (1), which develop following deployment and exposure in individuals in the absence of known prior respiratory illness or predisposition. Epidemiologic studies have previously associated exposure with emergent respiratory disease, both in the context of SW Asia and Afghanistan deployment (1, 2) and domestic air pollution (3). Additional studies of airborne exposures extend their impact beyond inflammatory and remodeling diseases of the respiratory tract and implicate these exposures in the development of lung cancer, cardiovascular disease and neuropsychiatric disease (4–7). This raises additional concerns about long-term sequelae of deployment exposures. However, our mechanistic understanding of the impacts of deployment exposures in relationship to disease pathogenesis and long-term risk remains fragmentary.

Previous studies have used cell and animal-based models of particulate matter (PM) exposure to identify potentially relevant cellular, physiologic and pathophysiologic responses. Although such studies fail to capture the complexity of true inhalational particulate exposure, they have been useful in defining PM-regulated pathways relevant to human exposures, such as Type I inflammation. For example, wood smoke particle (WSP) exposure resulted in increased inflammatory markers in the bronchoalveolar lavage (BAL) fluid of healthy human subjects (8) and induced pro-inflammatory gene expression in submerged airway epithelial cell culture (9). Building on these physiologic consistencies, other studies identified mediators that may contribute to airways diseases, such as mucus abnormalities (10) or cilia dysfunction (11), as well as long term sequelae, such as cellular dysplasia and malignancy (12). Additional studies connected inflammatory responses to particulate exposure with development of fibrosis (13) or allergic sensitization (14), underscoring the potential for particulate exposures to contribute to new-onset disease. More recently, particulate matter models specific to Afghanistan deployment (Afghan Desert Particulate matter or APM) have demonstrated physiologic responses relevant to DRRD, such as inflammation or airway hyperresponsiveness (15), and induction of cytokines such as CXCL8 and IL-33 (16). However, these studies did not identify the mechanistic basis for these responses, nor did they examine factors that may distinguish the effects of APM in comparison to other PMs that are associated with a different spectrum of disease risk.

Compounding the unknowns underlying particulate biology, the interactions between transcription factors that respond to external stimuli and the underlying lineage-specific transcriptional programming of the airway epithelia are poorly understood. We have previously described that multiple transcription factor programs are responsible for the acute transcriptional response to particulate exposure (17). To better conceptualize the interplay between transcription factors in response to exposures, we have adopted a classification paradigm (18) in which transcription factors are grouped based on their broad function in programming the cellular transcriptome. In this paradigm, lineage-determining transcription factors (LDTFs) establish and maintain cell-type specific chromatin architecture, which forms the genomic context in which subsequent signals enact transcriptional responses. Signal-dependent transcription factors (SDTFs) instead, but not mutually exclusively, direct transcriptional responses to specific stimuli. However, this activity is frequently determined by the architecture that arises from the activity of LDTFs (19). Although well established in other cell types (20), the interplay between LDTFs and SDTFs in the airway epithelia has not been characterized in the context of exposures. This mechanism is critical in understanding how exposures disrupt normal airway epithelia cell function.

Our overall objective in this study was to identify acute responses to a deployment-relevant exposure in small airway epithelial progenitor cells. This cell type features unique lineage-specific transcriptional programming (21) and is a critical cell type in the development of constrictive bronchiolitis. Although extrapolating acute responses to understand the development of a chronic, exposure-associated disease is challenging, we interpreted our results in the context of a multi-hit hypothesis, where acute responses mediate susceptibility to further insults that are ultimately compounded into a dysregulated disease state. We developed a comprehensive set of acute responses to APM exposure with both epigenetic and transcriptional effects by performing ATAC-seq and PRO-seq on SAECs. Through comparison with a non-deployment PM exposure, WSP, we identified responses unique to APM exposure. Finally, we identified genomic regions that shared dynamic responses in chromatin accessibility and nascent transcription to APM as candidate sites for interactions between SDTFs and LDTFs. Genome-wide analysis of these regions identified the P53/P63 family as enriched within this intersection and likely to play a significant role in intersecting small airway progenitor cell fate and function in response to APM exposure.

## Methods

### Cell culture

Cell culture of primary small airway epithelial cells (SAECs) was performed as described previously (22). Briefly, de-identified primary human small airway epithelial cells (< 2 mm diameter) were obtained from the National Jewish Health Biobank from donors with no history of deployment or chronic lung disease. Cells were first co-cultured with irradiated NIH/3T3 (ATCC) fibroblasts in F-medium containing 1 uM Y-27632 (APEX Bio) and then passaged onto tissue culture dishes double-coated with Type I bovine collagen solution (Advanced Biomatrix) and grown to confluence in BronchiaLife Epithelial Medium with the complete LifeFactors Supplement Kit (Lifeline Cell Technology). All exposures were performed in this medium. All cells were maintained in 5% CO_2_ at 37°C.

### Afghan Desert Particulate Matter – Preparation and Exposure

Afghan Desert Particulate Matter (APM) was a generous gift from Drs. Gregory Downey and Hong Wei Chu, who obtained this material from Drs. Brian Wong and Karen Mumy at the Naval Medical Research Unit (NAMRU) in Dayton, Ohio. The composition and cell culture exposure conditions of this material have been characterized previously (16). Briefly, topsoil from Bagram Air Force Base in the Parwan Province of Afghanistan was collected in August 2009 and shipped to the United States, where it was autoclaved and irradiated in accordance with Department of Agriculture standards. The material was then aerosolized into an exposure chamber using a Wright Dust Feeder (Mesa Labs, NJ) and size fractionated. Particle size was monitored using an optical sensor, gravimetric filter, and cascade impactor sampler. Settled particulates were collected from the bottom of the chamber and re-sterilized using gamma irradiation. Scanning electron microscopy determined that 89.5% of the particles were < 2.5 um in diameter, 10.4% between 2.5 and 10 um and 0.1% greater than 10 um. Dry APM was resuspended in 1X Phosphate buffered saline (PBS, Dulbecco) and vortexed prior to addition to cell culture media. After addition, media was gently agitated to ensure even distribution of the particles.

### Wood Smoke Particles – Preparation and Exposure

Wood Smoke Particles (WSP) were generously gifted by Dr. Andrew Ghio at the Environmental Protection Agency Laboratory in North Carolina and generated as described previously (8). WSP composition, preparation and exposure were described previously by our laboratory (22). The mean diameter of the freshly generated wood smoke particles was 0.14 um. After receipt of dry WSP, particles were suspended in 1X PBS and disaggregated by sonication and sterilized under UV light for 20 minutes. WSP particle suspension was then added directly to cell culture media at concentrations of 0.001 mg/ml to 1 mg/ml as indicated in the text and gently agitated to evenly disperse particles.

### Interleukin 8 Quantification by Enzyme linked Immunosorbent Assay (ELISA)

Quantification of IL-8 was performed by a commercial ELISA kit (R&D Systems). Briefly, plates were prepared with anti-IL-8 capture antibody prior to addition of sample. Cell supernatant was harvested following exposure and applied directly to prepared plates. Anti-IL-8 detection antibody was then added, and IL-8 levels were quantified from streptavidin-Horseradish peroxidase associated with the detection antibody using absorbance read by an Infinite M1000 plate reader (Tecan).

### Precision Run-on Sequencing (PRO-seq)

Cells required treatment and harvest on two separate days to generate enough nuclei to perform PRO-seq on two replicates per condition (untreated control (Ctrl) vs APM). For each set of replicates, Sm1 cells (small airway epithelial cells from donor 1) were seeded on 5 × 15 cm double collagen-coated plates per treatment and grown to confluence (3-4 days), then left untreated (Ctrl) or treated with APM (50 μg/cm^2^) for 20 hours. Cells were harvested and nuclei prepared as reported previously (23). PRO-seq libraries for all replicates were then prepared simultaneously by subjecting one aliquot of 1e7 nuclei/sample to 3-minute nuclear run-on reactions in the presence of Biotin-11-CTP (PerkinElmer) following our previously detailed protocol (24). Uniquely indexed libraries were pooled and sequenced on an Illumina NextSeq instrument using 75 bp single-end reads by the BioFrontiers Sequencing Facility at the University of Colorado Boulder.

### PRO-seq Computational Analysis

PRO-seq data were processed using a standardized Nextflow pipeline (https://github.com/Dowell-Lab/Nascent-Flow). A detailed quality control report including trimming, mapping, coverage, and complexity metrics is included in Supplemental File SR1. TDF coverage files output by the pipeline, normalized by reads per million mapped, were visualized using the Integrative Genomics Viewer (IGV) genome browser (v. 2.8.0). FStitch (v. 1.0) and Tfit (v. 1.0) were used to identify regions with bidirectional transcriptional activity as described (22). Reference Sequences for hg38 were downloaded from the UCSC genome browser (May 18, 2018). DESeq2 (v. 1.20.0, Bioconductor release v. 3.7) was used to determine differentially transcribed genes between control and APM exposure. Functional annotation of differentially transcribed genes was performed using DAVID v. 6.8 (25). For bidirectional comparisons, all predicted bidirectional Tfit calls were aggregated using mergeBed (argument -d 60) from BEDTools (v. 2.28.0) to generate an annotation file. Counts were then calculated for each sample using multicov (BEDTools v.2.28.0), and DESeq2 was used to identify differentially transcribed bidirectionals between treatments.

### Assay for Transposase Accessible Chromatin using Sequencing (ATAC-seq)

Sm1 cells were grown as described above for PRO-seq and aliquots of cells were taken from confluent 15 cm tissue culture dishes after vehicle (PBS) or APM treatment. Sm2 and Sm3 cells were grown to confluence in 6-well tissue culture dishes and treated with vehicle (PBS), APM or WSP for 20 hours. Cells were rinsed and scraped in ice-cold PBS, then ~50K cells from each treatment were pelleted and processed in duplicate for Omni-ATAC-seq using the protocol developed by Corces et al (26). Uniquely indexed libraries were pooled and sequenced on an Illumina NextSeq using 37 bp single- or paired-end reads by the BioFrontiers Sequencing Facility at the University of Colorado Boulder or on an Illumina NovaSeq using 150 bp paired-end reads at the University of Colorado Anschutz Medical Campus Shared Genomic Resource.

### ATAC-seq Computational Analysis

Computational analysis of ATAC-seq data was performed as described previously (22). Briefly, ATAC-seq reads were trimmed for adapters, minimum length, and minimum quality using the bbduk tool from the BBMap Suite (v. 38.73) with arguments ‘ref=adapters.fa ktrim=r qtrim=10 k=23 mink=11 hdist=1 maq=10 minlen=20’. Samples sequenced using 150 bp reads were further trimmed to 37 bp in length for consistency using the additional argument ‘ftr=36’. Trimmed reads were mapped to the human genome (hg38; downloaded from the UCSC genome browser on September 16, 2019, with corresponding hisat2 index files) using hisat2 (v. 2.1.0). Duplicate reads were removed using picard tools (v.2.26.6; http://broadinstitute.github.io/picard/) MarkDuplicates and deduplicated and sorted BAM files were converted to bedGraph coverage format using genomeCoverageBed from the BEDTools suite (v. 2.29.2). Read coverage was normalized to reads per million mapped using a custom python script and files converted to TDF format using igvtools (v. 2.5.3) for visualization in IGV. MACS2 (v. 2.1.4) callpeak with ‘--SPMR’ argument was applied to sorted BAM files for each replicate pair of samples with parameters set to call peaks with log_2_ fold change > 1 above baseline with q < 1e-5 as significant.

### Venn Diagrams for Feature Intersection

ATAC-seq peaks called by MACS2 and PRO-seq regulatory elements identified by Tfit were intersected using the BEDTools suite (v. 2.29.2) command “bedtools intersect.” Briefly, this operation took standard bed format input from two files and returned regions of the first file that had at least 1 base pair overlap with regions of the second file.

### Differential Transcription Factor Motif Enrichment

Transcription factor enrichment analysis for both PRO- and ATAC-seq datasets was performed using TFEA (v.1.0; https://github.com/Dowell-Lab/TFEA; (27)). Briefly, sets of consensus regions of interest (ROIs) are first defined for all Tfit-called bidirectionals (for PRO-seq) or all MACS2-called peaks (for ATAC-seq) using the muMerge function of TFEA. TFEA then calculates read coverage for each replicate within each ROI and applies DESeq2 to determine differentially represented ROIs by treatment condition and assign rank as a function of p-value. For each ranked ROI, the FIMO tool (28) from the MEME suite scans the 3 kb sequence surrounding the ROI center (bidirectional origin or ATAC-seq peak summit) for positions of transcription factor consensus motif matches (with p-value cutoff of 1e-5), represented by a curated set of position weight matrices (29). Enrichment scores (E-scores) for each motif are then calculated and corrected for sequence content to reduce known biases associated with local GC enrichment and p-values are determined using Z-scores. Motif displacement distributions are displayed as dot-plots where the location of all motif instances within ±1500 bp are indicated relative to the ROI center on the vertical axis, and the rank of each ROI based on signal loss or gain is indicated on the horizontal axis.

### Western Blotting

Western blotting and protein detection were performed using standard protocols (30) in SAECs cultured to confluence in 6-well plates and treated with vehicle control or APM. Primary antibodies purchased included anti-GRHL2 (Sigma HPA004820), anti-pan-TEAD (Cell Signaling Technologies 13295S) and anti-GAPDH (Santa Cruz sc-25778). Secondary antibody was ECL Donkey anti-Rabbit IgG, HRP-linked F(ab’)_2_ fragment (NA9340) from GE Healthcare/Amersham.

### RNA purification and quantitative reverse transcription-PCR (qRT-PCR)

RNA purification and qRT-PCR was performed as described previously (22). Briefly, SAECs were grown to confluence in 12-well tissue culture dishes and treated with vehicle (PBS) or APM for 20 hours. Cells were harvested in TRIzol (Life Technologies) and RNA purified by PureLink RNA Mini Kit (Life Technologies) prior to qRT-PCR, performed with normalization to RPL19 as previously detailed (23). Sequences of primers used for qRT-PCR analysis are provided in Supplemental Table ST1.

### Statistics

Statistical comparisons for IL-8 ELISA and qRT-PCR assays were made by 2-tailed t-test using the Bonferroni correction when appropriate. These analyses were conducted using statistical software embedded in Open Office (Apache OpenOffice v 4.1.7 http://www.openoffice.org/welcome/credits.html).

All High Throughput Sequencing Data is publicly available through the Gene Expression Omnibus (GEO) with accession GSE201152. Subseries GSE201149 contains 20 ATAC-seq libraries and subseries GSE201150 contains 4 PRO-seq libraries.

## Results

### Primary small airway epithelial cells (SAECs) secrete IL-8 in response to APM and WSP exposure in submerged culture

To establish conditions to compare APM with other clinically relevant PM exposures, we exposed SAECs to APM (50 μg/cm^2^ culture surface area) for 20 hours, based on a previous time course and dose response analysis (16). We demonstrated an increase in interleukin 8 (IL-8) secretion in cells from three healthy non-smoking donors, consistent with prior studies (16) (Figure 1A). We next sought to compare APM exposure with an additional particulate exposure model that resulted in similar inflammatory responses but had different clinical associations. We selected wood smoke particles (WSP), a highly relevant model for wildfire smoke exposure (31), and measured IL-8 secretion in response to varied doses of WSP exposure after 20 hours (Supplemental Figure SF1). Primary SAECs from the same three donors exhibited a similar increase in IL-8 secretion in response to APM exposure at 50 μg/cm^2^ and WSP exposure at 1 mg/ml (Figure 1B). Therefore, we selected these doses of APM (50 μg/cm^2^) and WSP (1 mg/ml) for subsequent genome-wide analysis and comparisons. Total particulate exposure volume was similar but different metrics were used based on convention.

**Figure 1.**
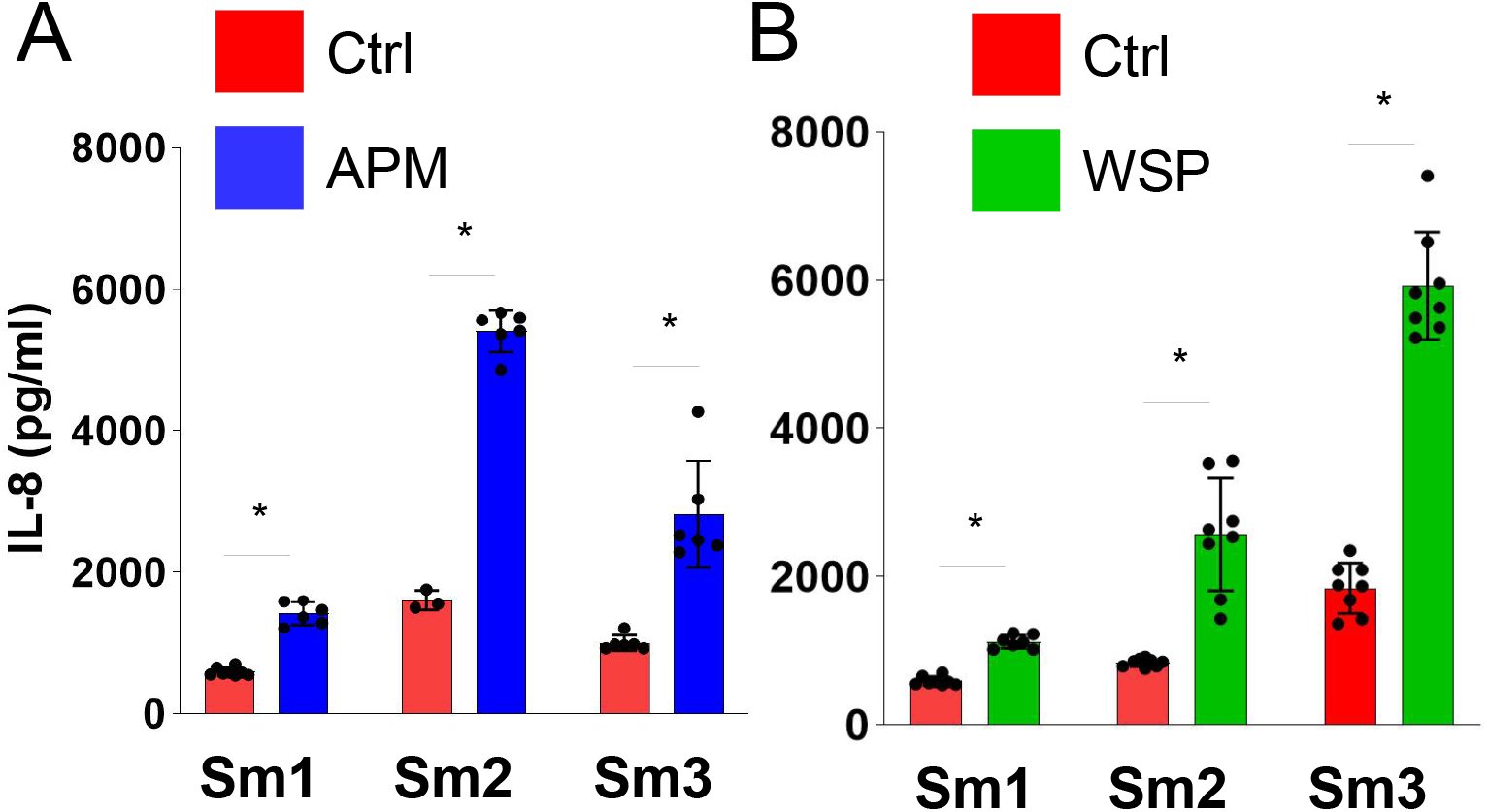
Particulate exposure induces IL-8 secretion in SAECs derived from three independent healthy donors. ELISA analysis of IL-8 concentration in the supernatant of Sm1, Sm2 and Sm3 SAECs exposed to (A) APM (50 ug/cm^2^ of well surface area) or (B) WSP (1 mg/ml of cell culture media) for 20 hours in submerged cell culture. Bars indicate mean IL-8 concentration (±SD); n ≤ 3/group, *p < 1e-3 for indicated comparisons.

### ATAC-seq (Assay for Transposase Accessible Chromatin with sequencing) identifies chromatin accessibility changes induced by APM exposure

Changes in chromatin architecture in response to a stimulus are generally mediated by the activity of transcription factors and the context of the underlying chromatin architecture of the particular cell type (32). To determine and compare regulation of chromatin changes in response to APM exposures, we performed ATAC-seq in SAECs following exposure to vehicle, APM or WSP. Principle Component Analysis (PCA) showed that the ATAC-seq data clusters predominantly by donor rather than exposure (Supplemental Figure SF2), reflecting the utility of ATAC-seq to identify basal differences in chromatin architecture in airway epithelial cells obtained from different donors. We next identified dynamic chromatin accessibility features in the three SAEC lines following APM exposure (Figure 2A) and compared the control- and APM-exposed ATAC-seq datasets using Transcription Factor Enrichment Analysis (TFEA).

**Figure 2.**
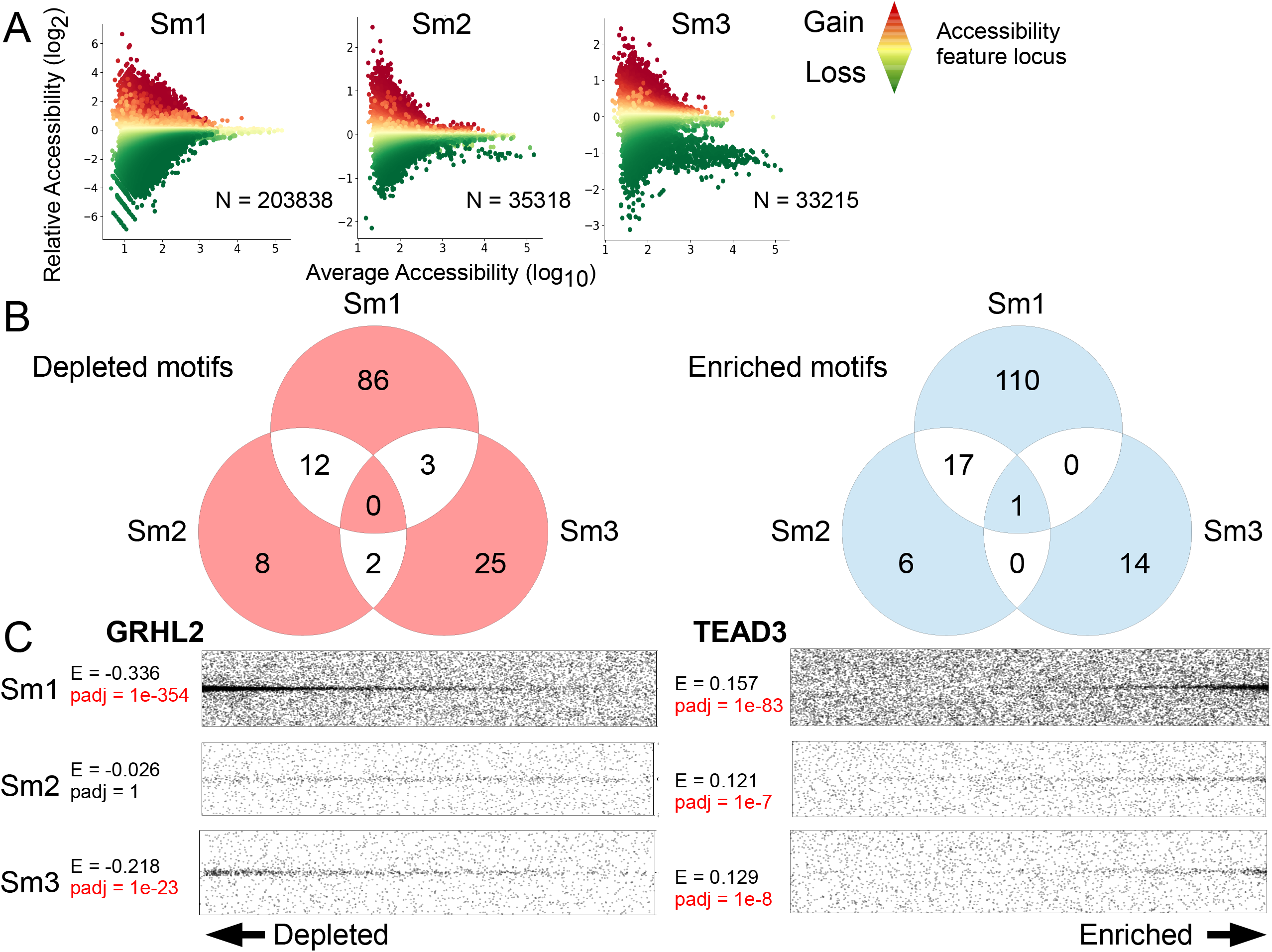
ATAC-seq identifies transcription factors associated with chromatin accessibility changes in response to APM exposure. (A) Pyramid plots illustrate MACS2-called ATAC-seq peaks (discrete dots) ranked by magnitude of accessibility gain (more red) or loss (more green) in response to APM exposure, as output during Transcription Factor Enrichment Analysis (TFEA). The total number (N) of unique loci for each SAEC line is listed below each plot. (B) Venn diagrams depict TFEA-identified numbers of transcription factor binding motifs for each SAEC line that were significantly (p_adj_ < 1e-5) depleted (motifs that cluster with ATAC-seq features that lose accessibility in response to APM) or enriched (motifs that cluster with ATAC-seq features that gain accessibility in response to APM) following APM exposure and their overlap between SAEC lines. (C) Representative dot plots for one depleted (GRHL2 – shared between Sm1 and Sm3, top right of Venn diagram in panel B) and one enriched (TEAD3 – shared between Sm1, Sm2 and Sm3, center of Venn diagram in panel B) motif identified by TFEA. Each dot represents the indicated motif encountered within ±1500 bp of an accessibility region, ranked by accessibility gain on the horizontal axis and distance to accessibility region center (with midpoint = 0) on the vertical axis. Enrichment (E) scores (a relative measure of motif enrichment between control vs APM exposure) and associated p_adj_ values are listed, with significant p_adj_ values indicated in red.

Significantly enriched or depleted transcription factor binding motifs (p_adj_ < 1e-5) were compared between SAEC lines (Figure 2B). Despite the divergent baseline chromatin accessibility profiles between the three donor-derived lines, concordant changes in chromatin accessibility for specific transcription factor binding motifs were identified across the three lines. A complete list of these transcription factors is provided in Supplemental Table ST2. In Figure 2C, these data are visualized for two specific shared motifs using dot plots, which depict the co-clustering of the GRHL2 motif with depleted chromatin accessibility features and the TEAD3 motif with enriched features, implying loss of GRHL2 and gain of TEAD3 activity. Using Western Blot, we confirmed expression of GRHL2 and TEAD3 in all three SAECs and demonstrated that the changes in factor binding motif enrichment or depletion described above were unrelated to expression changes (Supplemental Figure SF3).

### ATAC-seq distinguishes exposure- and donor-specific responses

To determine whether chromatin responses to particulate matter in SAECs are stereotyped or instead may be influenced by particulate characteristics, we performed ATAC-seq on Sm2 and Sm3 SAECs after 20 hours of WSP exposure and compared these responses to APM exposure responses. We identified a high degree of overlap between ATAC-seq peaks from vehicle- or WSP-treated samples and control- or APM-treated samples (Figure 3A). We again employed TFEA to identify transcription factor binding motifs associated with dynamic chromatin accessibility resulting from exposures and compared significant motifs (p_adj_ < 1e-5) between two donors and two exposures (Figure 3B). Shared motifs reflected stereotyped chromatin accessibility responses to exposure, but we also identified divergent motifs that represent exposure- or donor-specific responses. Complete TFEA results for all significant transcription factor binding motifs from all four samples are listed in Supplemental Table ST3. Depletion of P53/P63 motifs, which are extremely similar, was shared between both SAECs and both exposures, as demonstrated using dot plots (Figure 3C). In contrast to APM exposure, the TEAD3 motif was not enriched in either Sm2 or Sm3 in response to WSP exposure (Figure 3D). Thus, the SAEC response to particulate exposure may vary by donor and the underlying particulate composition, but TEAD3 activity is implicated as a distinguishing feature in chromatin responses to APM.

**Figure 3.**
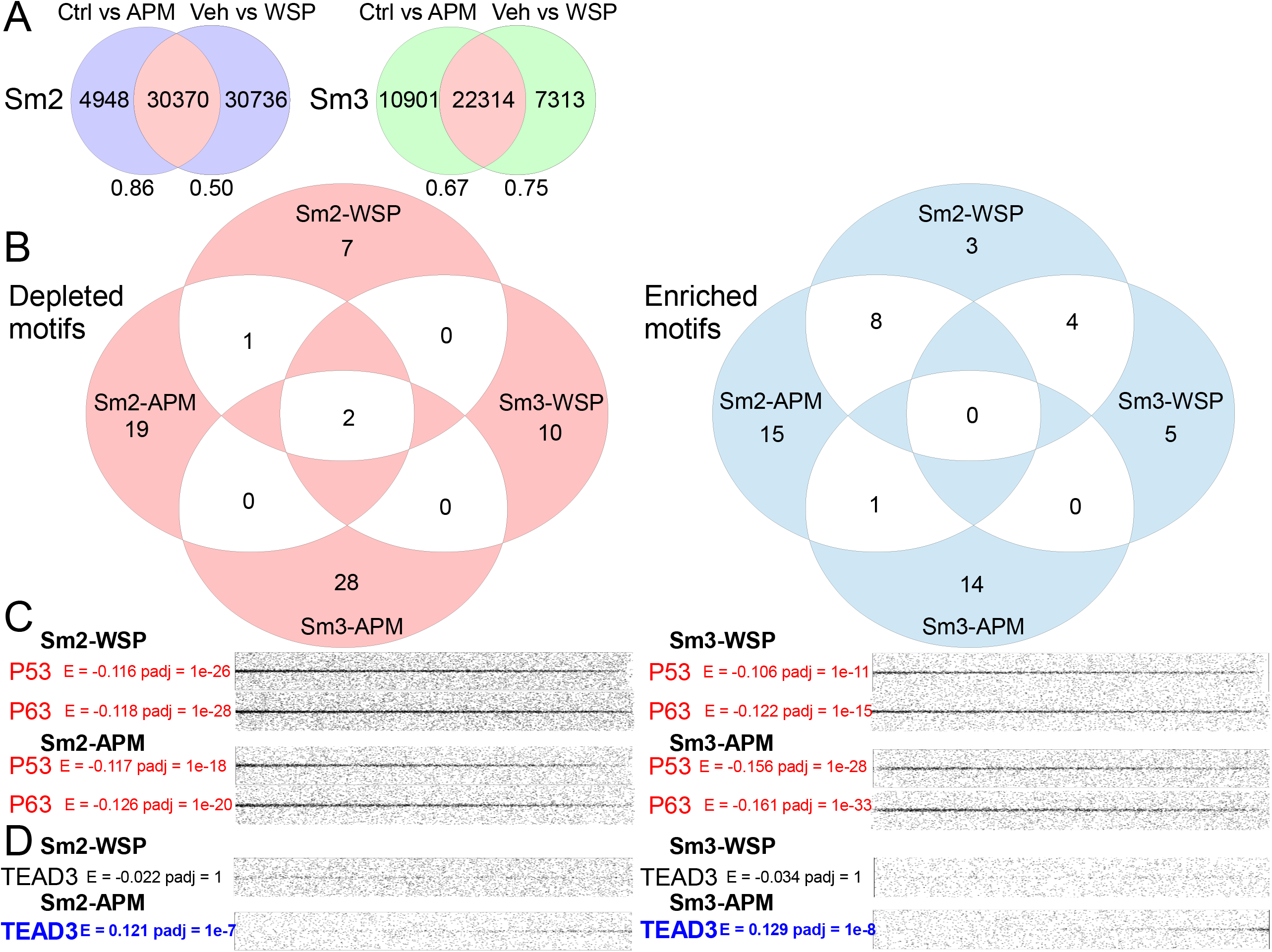
Chromatin accessibility profiles following APM vs WSP exposure distinguish exposure- and donor-specific responses. (A) Venn diagrams delineate overlap of MACS2-called ATAC-seq peaks identified in Sm2 or Sm3 cells exposed to APM vs WSP based on genomic coordinates. Numbers below Venn diagrams indicate the fraction of peaks from each indicated comparison found in the shared (pink) set. (B) Venn diagrams show TFEA-identified numbers of significantly (p_adj_ < 1e-5) enriched or depleted transcription factor binding motifs following APM or WSP exposure and their overlap, both by exposure and by donor, as described for Fig. 2B. (C) Dot plots for P53 and P63 binding motifs that shared depletion in Sm2 and Sm3 in response to both APM and WSP exposures; significant depletion indicated in red text. (D) Dot plots for the TEAD3 binding motif that was enriched in Sm2 and Sm3 exposed to APM but not WSP; significant enrichment indicated in blue text.

### Nascent transcript sequencing uncovers transcriptional pathways responsive to APM exposure

We next employed PRO-seq, which identifies nascent transcripts, such as gene transcripts and short-lived non-coding enhancer RNAs (eRNAs), with high temporal and spatial resolution, to examine the direct transcriptional response to APM exposure in SAECs. Our previous study demonstrated marked temporal variation in transcriptional responses within the first two hours of particulate exposure (22). Therefore, we anticipated that nascent transcription after 20 hours of exposure represents a steady-state transcriptional response to APM. We performed PRO-seq on a primary culture of human SAECs, Sm1, after 20 hours of APM exposure and used DESeq2 to identify gene transcription changes induced by APM. We identified 946 genes with increased transcription and 531 genes with decreased transcription in response to APM (Figure 4A). Representative PRO-seq genomic tracks visualized in the IGV Genome Browser for *GAS1* (downregulated) and *CXCL8* (upregulated) are demonstrated in Figure 4B. Gene transcription data from PRO-seq correlated with mRNA-qPCR from Sm2 and Sm3 cells exposed to APM for 20 hours (Supplemental Figure SF4). DAVID functional annotation (25) identified common pathways among APM-upregulated genes that included cell-cell adhesion (p = 0.0064) and keratin-associated signaling (p = 0.00013). Pathways shared among downregulated genes included cytochrome P450 (p = 2e-6) and glycolysis (p = 0.00064). The top 10 functional annotation pathways for downregulated and upregulated genes are depicted in Figure 4C. Thus, aligned with prior reports of particulate exposures, APM exposure causes significant changes in gene expression associated with specific biologic processes.

**Figure 4.**
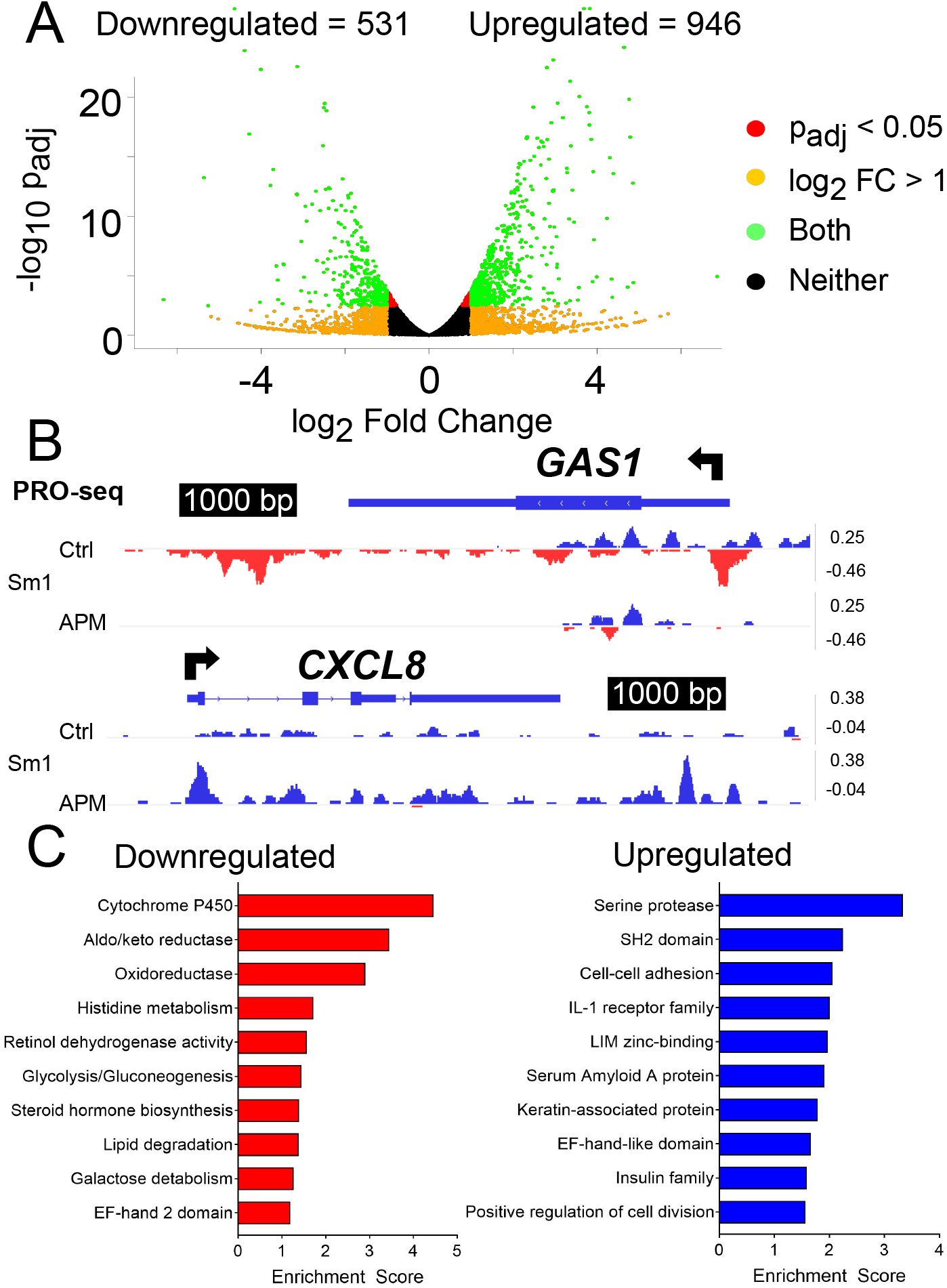
PRO-seq reveals sustained transcriptional programs associated with APM exposure. (A) Volcano plot illustrates differentially regulated nascent gene transcripts in Sm1 cells treated with APM for 20 hours. Each point represents one gene, with red indicating p_adj_ < 0.05, light orange representing log_2_ fold change > 1, green indicating both p_adj_ < 0.05 and log_2_ fold change > 1 and black indicating neither condition was satisfied. The number of genes meeting both criteria (green dots) that were down- or upregulated is shown. (B) Representative examples of downregulated (left) and upregulated (right) genes shown as PRO-seq tracks visualized in the Integrative Genomics Viewer (IGV) genome browser based on counts per million mapped reads (vertical scales). Positive (blue) indicates reads annotated to the sense strand while negative (red) data reflect reads annotated to the antisense strand. The TSS and direction of transcription are indicated by arrows at the top of each panel. (C) Bar graphs display top 10 most significantly enriched functional annotation terms output by DAVID Functional Annotation Clustering applied to downregulated (left) and upregulated (right) transcripts.

### Nascent enhancer RNA transcripts identify transcriptional regulatory elements and transcription factors that mediate responses to APM

It is increasingly recognized that SDTFs integrate the cellular programming of lineage-determining factors to exact cell-type specific responses to stimuli, such as inflammation or exposures (33). To identify SDTFs underpinning changes in gene expression caused by APM in small airway epithelia, we quantitated Transcriptional Regulatory Elements (TREs) using Transcription-fit (Tfit) (34). Tfit interrogates PRO-seq transcripts for eRNA signatures, which demarcate TREs, and TRE activity is estimated by quantification of eRNA transcripts at the specific locus. Differential expression analysis of 20,190 TREs identified 207 TREs with significantly increased eRNA transcription (p_adj_ < 0.05) and 310 TREs with decreased eRNA transcription (Figure 5A). TFEA identified a series of depleted and enriched transcription factor binding motifs associated with transcriptional changes at TREs (Figure 5B). We next defined the intersection between PRO-seq TREs from APM-exposed Sm1 cells and ATAC-seq features from APM-exposed SAECs based on genomic coordinates (Figure 5C). Simple Enrichment Analysis (35) identified common motif occurrences within each set of intersected regions (Supplemental Figure S4F). Representative genome tracks in Figures 5D-E show differentially transcribed TREs neighboring *MAB21L4* and *AKR1C1*. These regions also exhibit dynamic chromatin accessibility in select SAEC lines, suggesting that transcription factors mediating APM exposure responses at these loci exhibit both signal-transduction and lineage-determining properties.

**Figure 5.**
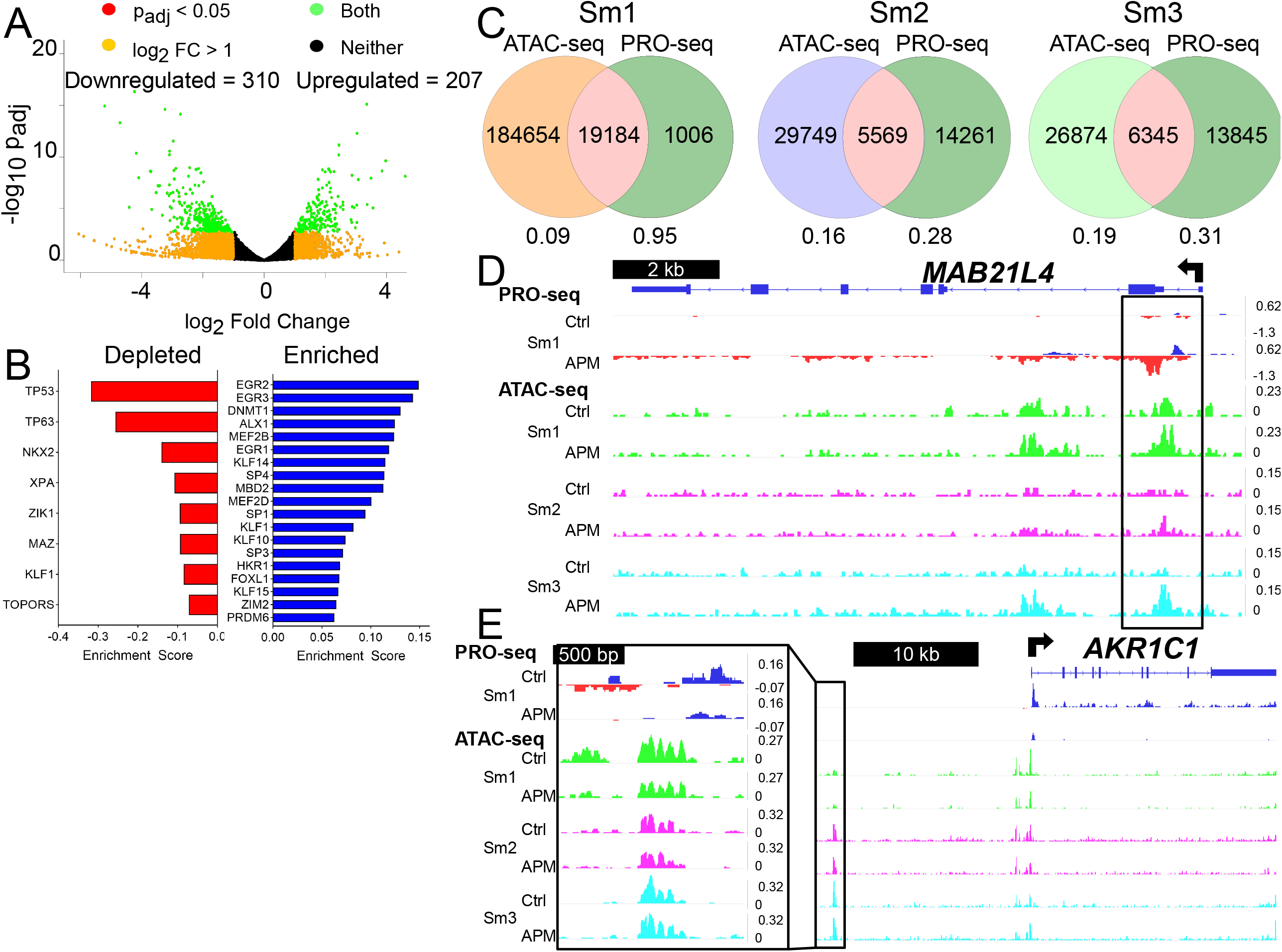
TREs defined by PRO-seq bidirectional signatures identify a distinct set of transcription factors that respond to APM exposure. (A) Volcano plot illustrates differentially regulated Transcriptional Regulatory Element (TRE) transcripts in Sm1 cells treated with APM for 20 hours, as described for Fig. 4A. (B) Transcription factor binding motifs identified by TFEA within PRO-seq data are represented as bar graphs. In this case, depleted motifs refers to those encountered more frequently in TREs with loss of transcriptional signal, while enriched motifs are those encountered more frequently in TREs with gain in transcriptional signal. Motifs listed meet the threshold of p_adj_ < 1e-5 and are ranked by E score, here a relative measure of motif enrichment within TREs. (C) Venn diagrams depict the total numbers of control and APM-exposed ATAC-seq peaks from each SAEC line (ATAC-seq), control and APM-exposed PRO-seq TREs from Sm1 cells (PRO-seq), and their overlap, based on genomic coordinates. Numbers below each diagram represent the fraction of ATAC-seq or PRO-seq peaks found within the intersection for each SAEC (pink). Representative examples of one upregulated TRE near the *MAB21L4* promoter (D) and one downregulated TRE 5’ to the *AKR1C1* gene locus (E) shown as PRO-seq tracks from Sm1 exposed to APM and ATAC-seq tracks from three SAEC lines exposed to APM, as visualized in IGV. ATAC-seq tracks are color-coded by SAEC line, with green = Sm1, purple = Sm2, and cyan = Sm3. Vertical scales on all tracks indicate counts per million mapped reads.

### Integrating genomics approaches delineate signal-transduction and lineage-determining transcription factor mediators of APM exposure

We reasoned that regions of both dynamic transcription and chromatin accessibility in response to APM exposure in small airway epithelial progenitor cells likely represent cooperative activity of SDTFs and LDTFs. To interrogate these regions genome-wide, we re-applied TFEA to ATAC-seq data from three SAEC lines within TREs called from PRO-seq data. A complete list of significant motifs (p_adj_ < 1e-5) is provided in Supplemental table ST4. Using this approach, we again examined the TEAD3 and P53/P63 motif changes previously identified in our study.

The TEAD3 motif was not enriched in SAEC ATAC-seq analysis using PRO-seq annotations or Sm1 PRO-seq (Figure 6A). However, Sm2 and Sm3 cells exhibited increased gene expression of four canonical TEAD3-target genes following APM exposure, including *CCN1, CCN2, MYC* and *FGFBP1* (Figure 6B). Figure 6C demonstrates one example where, despite a significant increase in *CCN1* gene transcription, neighboring genomic features containing canonical TEAD3 binding sequences (JASPAR (36)) exhibit increased chromatin accessibility without changes in eRNA transcription. This suggests that TEAD3 does not directly affect transcriptional regulatory activity in the manner of a SDTF. Instead, TEAD3, in response to APM exposure, changes the underlying chromatin landscape to promote expression of target genes, either through cooperation with an unknown factor or inducing TRE-promoter looping, a function more characteristic of a LDTF (19).

**Figure 6.**
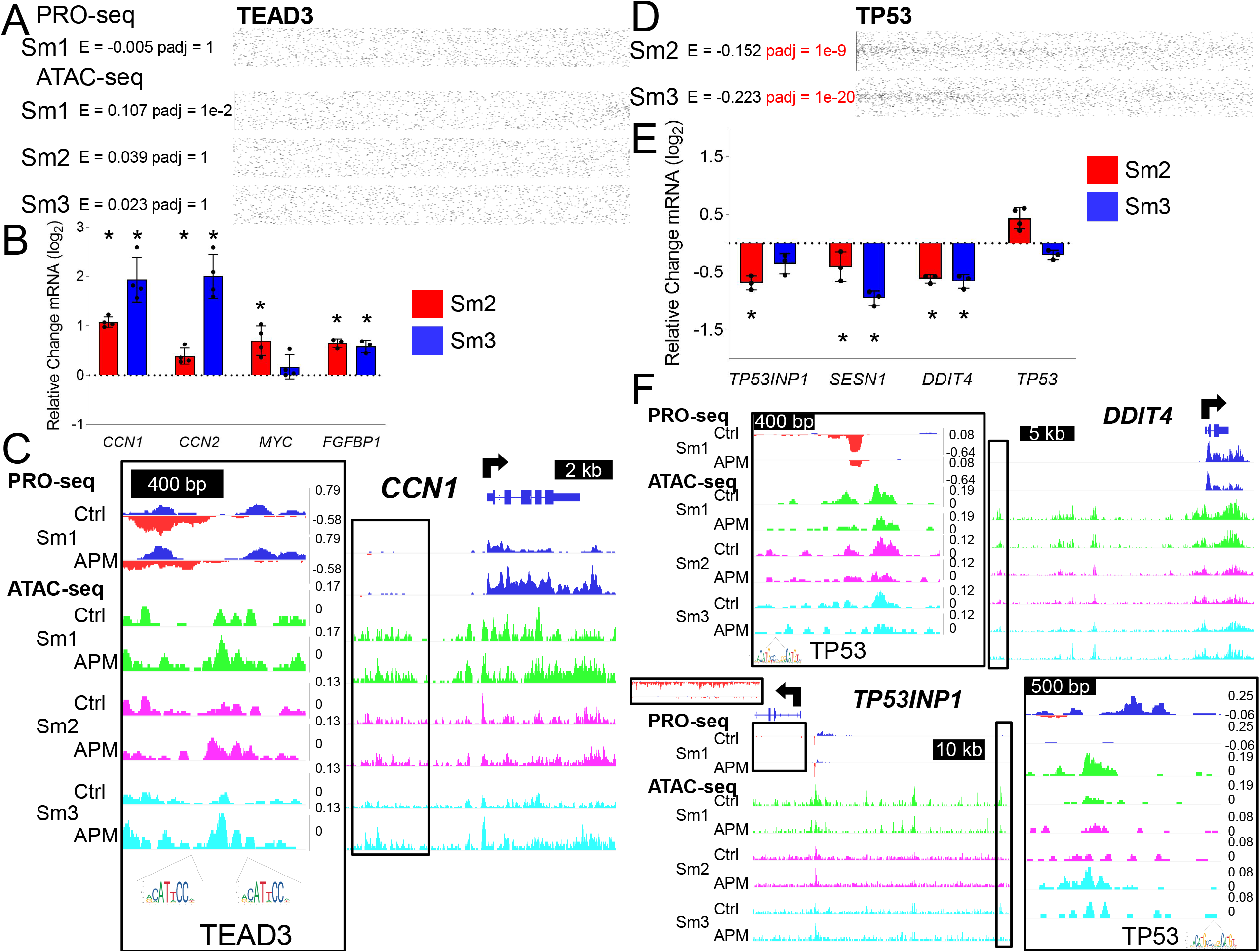
APM-responsive transcription factors exhibit different classes of transcriptional regulation. (A) Dot plots demonstrate no TEAD3 binding motif enrichment when PRO-seq-defined TREs are used to define regions for TFEA analysis of control vs APM-exposed SAECs. (B) qRT-PCR analysis of indicated gene expression in Sm2 or Sm3 cells following APM exposure for canonical TEAD targets *CCN1, CCN2, MYC* and *FGFBP1*. Bars represent mean normalized C_T_ values on a log_2_ scale (±SD) relative to vehicle-treated controls; n = 4/group, *p < 0.05 vs. vehicle. (C) IGV-visualized tracks of Sm1 control- and APM-exposed PRO-seq samples and three SAEC control- and APM-exposed ATAC-seq samples for one dynamic chromatin accessibility region 5’ to *CCN1* (boxed in black and magnified). Specific regions containing canonical TEAD3 binding sequences (JASPAR) are indicated as motif logos under the genome tracks. (D) TFEA analysis of control vs APM-exposed Sm2 and Sm3 ATAC-seq data within PRO-seq defined regions confirms significant (p_adj_ < 1e-5) depletion of the P53 binding motif, illustrated by dot plots. (E) qRT-PCR analysis of indicated gene expression in Sm2 or Sm3 cells following APM exposure for canonical P53 targets *TP53INP1, SESN1* and *DDIT4*; n = 4/group, *p < 0.05 vs. vehicle. (F) IGV-visualized tracks of Sm1 control- and APM-exposed PRO-seq samples and three SAEC control- and APM-exposed ATAC-seq samples for one TRE 5’ to the *DDIT4* gene locus and one TRE 5’ to the *TP53INP1* gene locus (boxed in black and magnified). Both TREs contain canonical P53 binding sequences (JASPAR) as indicated.

In contrast to TEAD3, we confirmed depletion of the P53/P63 motif in APM-exposed Sm2 and Sm3 ATAC-seq samples when examined only within TREs identified by PRO-seq (from Sm1 cells) (Figure 6D). Depletion of the P53 motif was corroborated by a decrease in gene expression of three canonical P53 target genes, *TP53INP1, SESN1* and *DDIT4*, following APM exposure (Figure 6E). TREs regulating *DDIT4* and *TP53INP1*, with regulation suggested by concordant transcriptional kinetics, contained canonical P53 binding sequences (JASPAR) and demonstrated simultaneous transcriptional and chromatin accessibility changes in response to APM exposure (Figure 6F). These data suggest a combined signal-transduction and lineage-determining role for TREs harboring P53/P63 motifs, and strongly implicate a role of P53/P63 in response to APM in airway epithelia.

## Discussion

The mechanistic basis for the development of deployment- and exposure-associated respiratory disease is poorly understood. Here, we approached this problem by exposing primary SAECs to APM, a highly relevant deployment-specific exposure, and characterizing the genomic responses. We identified several pathways that may be involved in the development of DRRD and potential neoplastic implications of deployment exposures, such as loss of P53/P63, GRHL2 or cytochrome P450 signaling. Furthermore, we identified transcription factors that mediate transcriptional responses, such as the EGR family, factors that affect chromatin accessibility change, such as the TEAD family, or factors with shared function, such as the P53/P63 family. We identified pathways specific to APM exposure by comparing effects of APM and WSP exposure, with TEAD activation, a pathway that previously had not been associated with particulate exposures, as specific to APM. In aggregate, our data indicate that APM exposure in human airway epithelial cells causes broad changes in transcription factor activity, with certain effects specific to APM in comparison to WSP.

The combined application of PRO-seq and ATAC-seq allowed us to partition responses to APM exposure into two categories. PRO-seq identifies direct transcriptional responses to a stimulus while ATAC-seq profiles changes in chromatin architecture. Although both responses are important to cell fate and function, we hypothesize that transcription factors that mediate these changes fall under two distinct but overlapping categories. LDTFs regulate the underlying chromatin architecture of the cell and, dependent on cell type or memory from previous stimuli, establish the chromatin context for subsequent exposures (20). We interpreted responses detected in ATAC-seq profiles to reflect the function of LDTFs. SDTFs directly control transcriptional responses within the context of chromatin architecture established by LDTFs (33). We interpreted PRO-seq data to reflect the functions of SDTFs. Using TFEA, we identified transcription factors from ATAC-seq and PRO-seq data and empirically classified them under these two categories.

We identified important SDTF pathways in SAECs exposed to APM that affect disease susceptibility to additional insults and support a multi-hit hypothesis for the development of DRRD. For example, APM exposure resulted in downregulation of cytochrome P450 signaling, which is responsible for catabolism of polyaromatic compounds (37). Even transient loss of this program can result in increased susceptibility to additional environmental exposures, as increased polyaromatic hydrocarbon composition in particulate exposures is associated with increased cytotoxicity (38) and several polyaromatic compounds are known carcinogens (39). Thus, a loss of cytochrome activity may predispose to aberrant responses and additional injury in response to repetitive exposures, which are typically a feature of DRRD.

Our data indicate that APM-mediated activation of the TEAD family of transcription factors is a unique feature of APM that relates directly to small airway epithelia lineage. In contrast to APM, WSP exposure did not result in TEAD enrichment, and with APM-exposure, TEAD enrichment was demonstrated only in dynamic chromatin accessibility features identified by ATAC-seq and not TREs identified by PRO-seq. TEAD transcription factors are classically identified as downstream effectors of the Hippo-YAP-TAZ signal transduction pathway that promote development and cell proliferation (40). Small molecules have been described that can activate or block Hippo pathway signaling (41, 42), and the TEAD family may represent a potential therapeutic target for the treatment of DRRD or mitigation of toxic responses to deployment-related exposures. Ultimately, further investigation is necessary to characterize the impact of TEAD-associated chromatin accessibility gain on subsequent exposures or cell fate.

P53/P63 signaling responses to APM were identified by interrogation of dynamic TREs from PRO-seq and chromatin accessibility features from ATAC-seq, suggesting a combined signal-dependent and lineage-determining role for P53/P63 in small airway epithelia progenitor cells. This latter function aligns with numerous previous studies of the airway epithelia (43, 44, 46, 48), however, the integrated role of this family in exposure responses has not been reported to our knowledge. The combined role may reflect differential activity of distinct P53/P63 family members or the effects of transcription factors that cooperate with P53/P63. For example, GRHL2, another LDTF identified by our study, has been demonstrated to work with P53/P63 in determining epithelial cell fate at the epigenetic level (45).

The importance of P53/P63 signaling in APM exposure is further corroborated by a series of studies that connect dysregulations in this family with bronchiolitis and small airways disease. Reduced P53 activity has previously been implicated in distal bronchiolar pathology and a failure of apoptosis in response to RSV infection (47), a common cause of acute bronchiolitis. Moreover, P63 expression is reduced in patients with constrictive bronchiolitis arising from mustard gas exposure (49). Although additional studies are needed to establish a causal relationship, our work suggests that reduced P53/P63 activity may predispose to bronchiolitis in the setting of APM exposure, either acting independently with multiple transcription factor roles or in association with additional factors, such as GRHL2. Further investigation of dynamic P53/P63 motif-containing TREs and chromatin accessibility features identified in our study may delineate the specific factors responsible for these multiple P53/P63 roles and determine how signal-dependent responses integrate with lineage factors in determining airway epithelia response to exposures. Finally, increasing P53 activity via agents such as idasanutlin (50), which is currently in clinical development for myeloid cancers (51), is feasible and may provide another therapeutic avenue in DRRD.

The study of exposure biology has multiple layers of complexity, which contribute to the poor understanding of DRRD. Factors such as concentration or duration of exposure are highly variable in patients with DRRD and difficult to model. Even in model system studies such as ours, the partition between homeostatic and pathophysiologic responses to exposure is not clear. However, ATAC-seq is one genomic method that, given the low input cell number, can be directly applied to patient-derived samples. In this study, we explored the utility of ATAC-seq to identify mechanisms of APM exposure and identified the TEAD family of transcription factors, a novel pathway not previously associated with particulate exposure. Furthermore, we demonstrated that the intersection of ATAC-seq data from multiple SAEC lines with TREs identified from a single PRO-seq experiment delineates a subclass of LDTFs that are shared between SAECs. Future studies that apply ATAC-seq to patient samples, such as small airway epithelial cells from distal bronchoscopic brush biopsy, can explore dysregulation in these core lineage factors by integrating those data with the TRE set identified by PRO-seq in this study. In addition, we used this set of TREs to investigate dynamic chromatin accessibility in multiple SAEC lines. Through this approach, we were able to delineate which transcription factors identified by ATAC-seq were consistently regulated by APM exposure or exhibited signal-dependent properties. Thus, future studies applying ATAC-seq to a variety of dynamic samples can use annotations from prior PRO-seq studies to classify transcription factors identified by the analysis. Given the high input requirements for PRO-seq, such as large cell numbers, time and sequencing cost, the annotation approach provides an easy-to-implement option when patient or exposure sampling methods preclude paired PRO-seq and ATAC-seq library preparation.

Although our study was not specifically powered for detecting individual-level variations in exposure responses, we demonstrated variation in genomic response patterns between SAECs from different donors. For example, GRHL2 loss was present in Sm1 and Sm3 but not Sm2. This variation may depend on several factors, such as age, sex, systemic disease or exposure history. Future studies that incorporate donor-level data can investigate the effect of systemic factors or genetic variants on responses to APM exposure. Finally, our approach also detected differences between the genomic responses to APM versus WSP exposures. It is tempting to speculate that these differences in our reductionist model portend the well-known clinical differences in response to deployment-associated particulates and other more common sources of PM.

Our model system had several limitations. SAECs were cultured and exposed in submerged cell culture without differentiation into a cuboidal ciliated epithelium, which is characteristic of small airways (52). The dose and composition of exposures may also diverge between our model and true inhalational exposures experienced by military personnel deployed to Southwest Asia. However, despite these limitations, our study identified several plausible and novel features of APM exposure that could only be identified through the use of sophisticated genomics tools in our model system. Extending our findings to *in vivo* models and even more sophisticated *in vitro* systems, such as precision cut lung slices, will be crucial for validating the novel APM-associated pathways we have reported here.

## Supporting information

Supplemental Items Legend

Supplemental Reports SR1

## Supplemental Items are available at

https://figshare.com/projects/Integrated_Genomics_Approaches_Identify_Mediators_of_Transcri_ptional_and_Epigenetic_Responses_to_Afghan_Desert_Particulate_Matter_in_Human_Small_Airway_Epithelial_Cells/138039

