## Supplementary figures and images for "Integrated Genomics Approaches Identify Transcriptional Mediators and Epigenetic Responses to Afghan Desert Particulate Matter in Small Airway Epithelial Cells"

### Supplemental Figure SF1

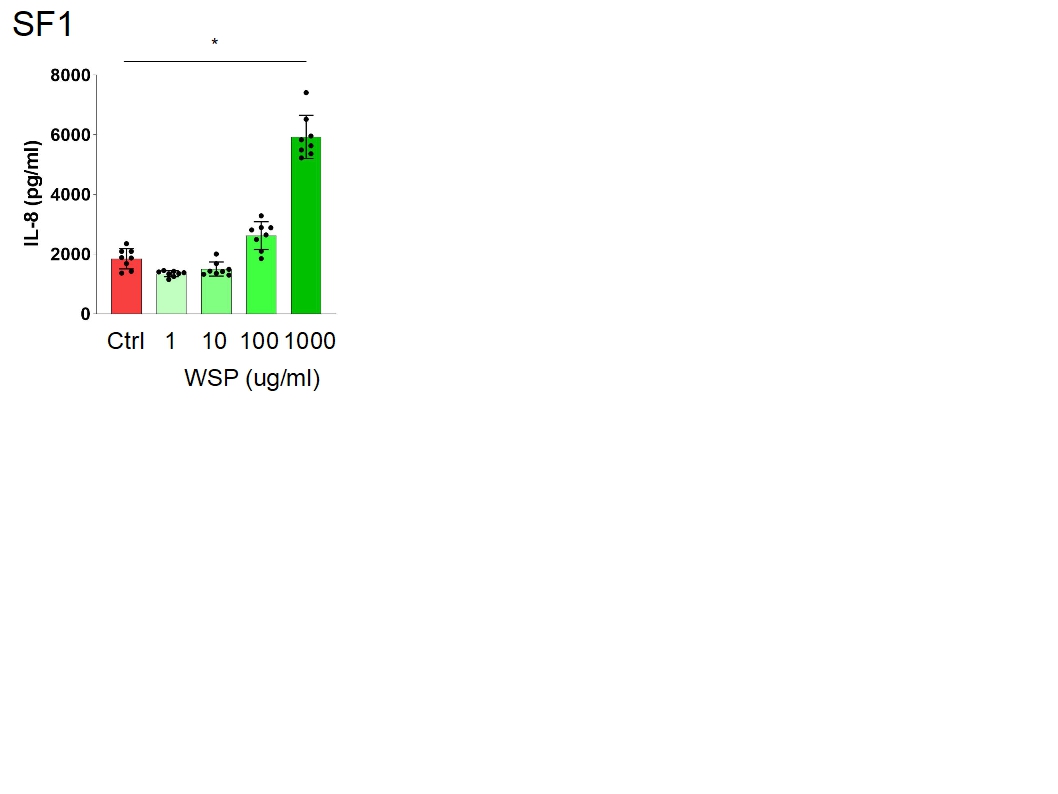

### Supplemental Figure SF2

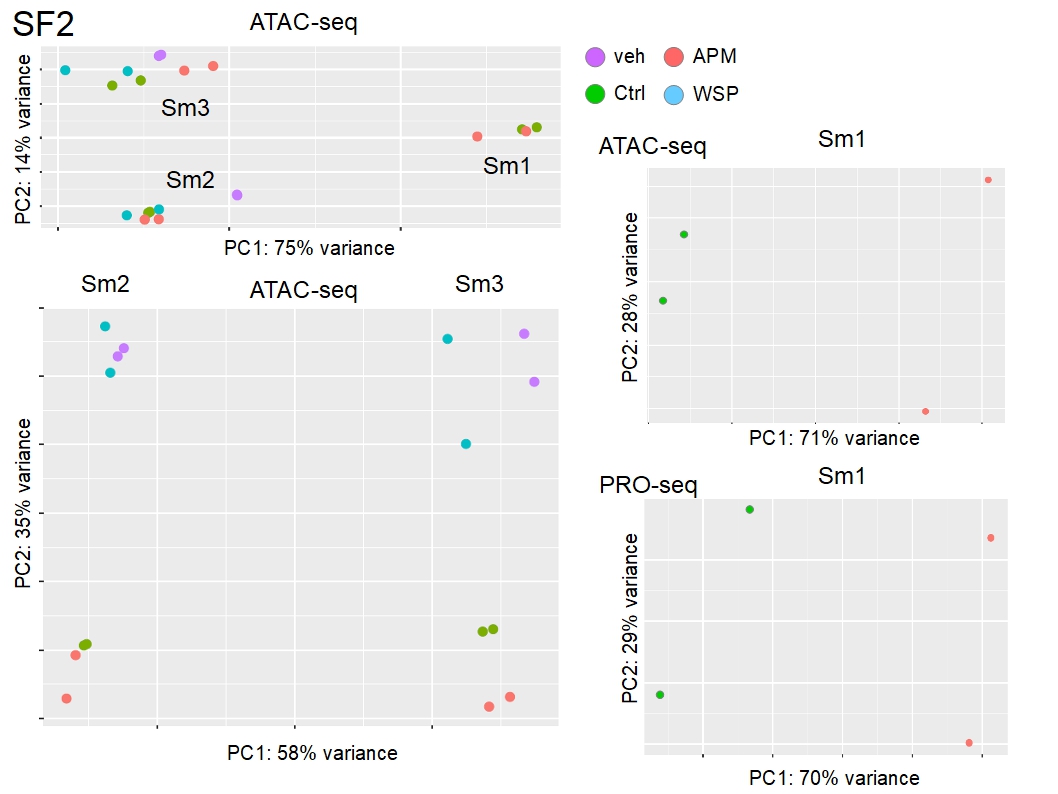

### Supplemental Figure SF3

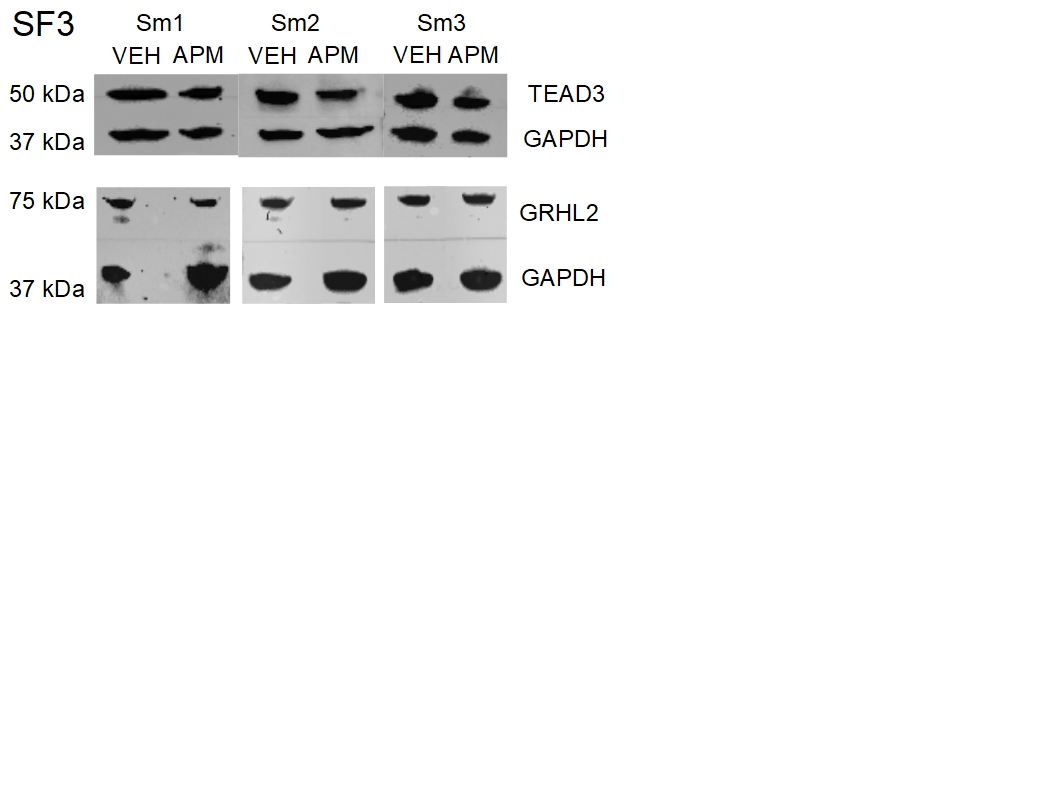

### Supplemental Figure SF4

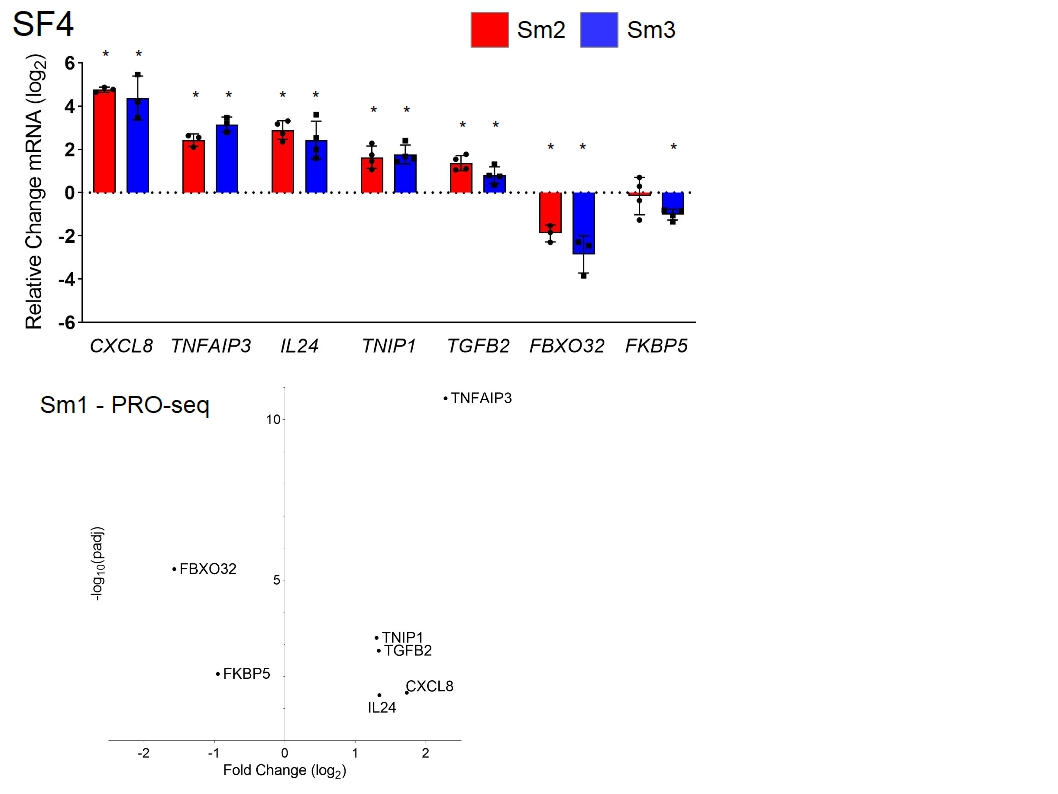

### Supplemental Figure SF5

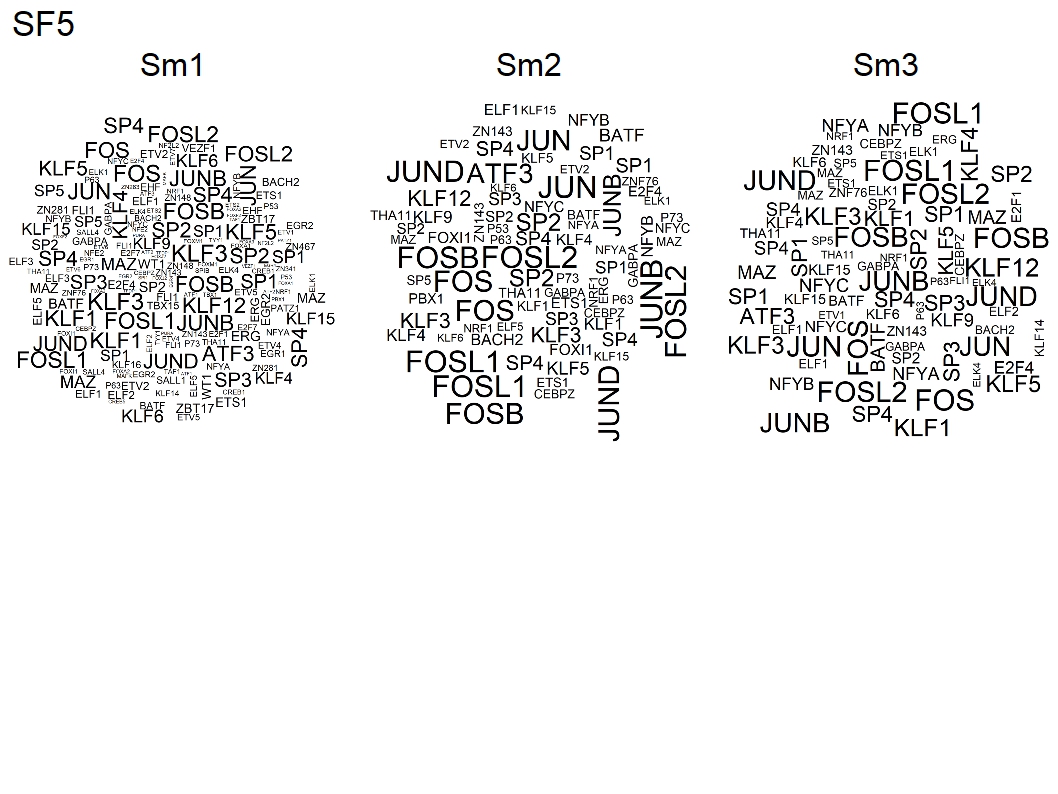
