## Supplemental Items Legend for "Integrated Genomics Approaches Identify Transcriptional Mediators and Epigenetic Responses to Afghan Desert Particulate Matter in Small Airway Epithelial Cells"

**Supplemental Items Legends**

Figure SF1 – IL-8 secretion dose response experiment of WSP exposure in Sm3. ELISA analysis of IL-8 concentration in the supernatant of Sm3 SAECs exposed to WSP at varied concentrations in cell culture media for 20 hours. Bars indicate mean IL-8 concentration (±SD); n = 6/group, *p < 1e-3 for indicated comparisons.

Figure SF2 – Principal Component Analysis plots compare ATAC-seq and PRO-seq sequencing data. Top left – PCA plot showing all 20 ATAC-seq samples from three SAECs, exposed to vehicle/control, WSP or APM. Bottom left – zoom in on Sm2 and Sm3 samples exposed to vehicle/control, WSP or APM. Note – since WSP and APM experiments were conducted independently, separate vehicle/control samples were used for baseline comparisons. Right – Sm1 ATAC-seq (top) and PRO-seq (bottom) PCA plots showing separation of samples by control or APM exposure.

Figure SF3 – Western blot demonstrates baseline expression of TEAD3 and GRHL2 without any change in factor expression following exposure to APM. GAPDH served as a loading control.

Figure SF4 – Top – qRT-PCR results of Sm2 and Sm3 cells exposed to APM for 20 hours for select target genes. Bars represent mean normalized CT values on a log2 scale (+SD) relative to vehicle-treated controls; n = 4/group, *p < 0.05 vs. vehicle. Bottom – PRO-seq gene transcript data analyzed by DESeq2 for select target genes. Points represent a normalized comparison between control and APM exposed samples in duplicate meeting the significance values of p_adj_ < 0.05 and |log_2_ fold change| > 1.

Figure SF5 – Word clouds represent motif enrichment within overlap genomic regions (Figure 5C) between macs2-called ATAC-seq peaks from each SAEC line exposed to APM or control and Tfit-called PRO-seq transcriptional regulatory elements from Sm1 cells exposed to APM or control. Word size is directly proportional to -log_10_(p_adj_) for enrichment with threshold p_adj_ < 1e-10.

Reports SR1 – MultiQC (quality control) report. A detailed quality control report of raw and processed PRO-seq data.

Table ST1 - qRT-PCR primer sequences used to quantify total RNA. *RPL19* was used as the reference.

Table ST2 – Significantly enriched and depleted TFEA motifs (p_adj_ < 1e-5) from SAEC-ATAC-seq analysis following APM exposure for all three SAECs, categorized by shared motifs to reflect the Venn diagrams in Figure 2B.

Table ST3 – Significantly enriched and depleted TFEA motifs (p_adj_ < 1e-5) from SAEC-ATAC-seq analysis for Sm2 and Sm3 cells exposed to APM or WSP, categorized by shared motifs to reflect the Venn diagrams in Figure 3B.

Table ST4 – Significantly enriched and depleted TFEA motifs (p_adj_ < 1e-5) from SAEC-ATAC-seq analysis for all SAEC lines exposed to APM within regions defined by all PRO-seq TREs in Sm1 cells.
